# Discriminating rare disease cases from their controls based on observed and excluded phenotypes

**DOI:** 10.64898/2026.09.15.751721

**Authors:** Yanting Guo, Yang Yao, Yiyao Zhang, Guangyou Duan, Ning Zhang, Shan Gao

## Abstract

Rare diseases are individually uncommon but collectively prevalent. Their primary clinical challenge lies not in treatment but in diagnosis. In the early stages of clinical management, it is frequently unclear whether the observed phenotypes are associated with a rare disease. Leveraging machine learning methods to mine latent associations between these phenotypes and rare diseases for early diagnosis offers a viable strategy to alleviate this diagnostic dilemma. In the present study, we demonstrated that machine learning can effectively discriminate rare disease cases from their controls using observed and excluded phenotypes as features. Among them, the Random Forest model achieved the best classification performance with a certain degree of generalizability. Based on further analysis of the feature selection results, we conclude that the two factors, specificity and occurrence count, are important for phenotype selection in rare disease discrimination, and comparable importance should be attached to both observed and excluded phenotypes, during feature construction.

## Introduction

Rare diseases are individually uncommon but collectively prevalent. According to an analysis of 6,172 unique rare diseases in Orphanet database [1], the evidence-based estimate of the global population prevalence of rare diseases ranges from 3.5% to 5.9%, which equates to 263–446 million persons affected worldwide at any point in time [2]. Their primary clinical challenge lies not in treatment but in diagnosis, known as the “diagnostic odyssey” [3]. Surveys indicate the average diagnostic delay is measured in years, during which patients are managed as if they had common diseases and undergo multiple referrals before a rare disease is even considered. Notably, such delays stem not primarily from disease complexity, but from the absence, during the initial evaluation, of both phenotypes and genetic or chromosomal evidence closely linked to a rare disease. In the early stages of clinical management, more often than not, whether the observed phenotypes are associated with a rare disease is unknown, offering little directional guidance for diagnosis. Leveraging machine learning methods to mine latent associations between these phenotypes and rare diseases offers a viable strategy to alleviate this diagnostic dilemma.

Constrained by data scarcity and low standardization, early machine learning-based research on rare diseases focused primarily on natural language processing and text mining techniques to accurately extract clinical phenotypes and genomic features of rare diseases from massive medical literature and databases. Subsequent research increasingly prioritized the adoption of emerging machine learning methods. For example, Alsentzer et al. introduced SHEPHERD [4], pioneering the first deep learning model tailored for personalized diagnosis in rare genetic disorders. However, the bedrock for the advances in this field was laid by the standardization of phenotypic data, exemplified most notably by the Human Phenotype Ontology (HPO) [5], as well as early phenotype-matching tools based on HPO, such as Phenomizer. Another notable contribution is Orphanet [1], the world’s most authoritative multilingual knowledge base for rare diseases and orphan drugs. Featuring its own classification system, Orphanet assigns a unique ORPHA code to each rare disease, thereby addressing the limitations of ICD-10 (International Statistical Classification of Diseases and Related Health Problems 10th Revision) in rare disease granularity—a framework subsequently deeply integrated into ICD-11 [6]. Despite these efforts, clear cross-linkage between disparate standardized sources is still wanting, with substantial clinical data remaining fragmented in literature or private databases, highlighting an urgent imperative for wide-scale standardization and integration. Phenopacket Store [7] is a milestone in integrating disparate standardized resources. This open-source database and software toolkit seamlessly integrates HPO and Orphanet by mapping rare diseases to ORPHA codes or the Mondo Disease Ontology (MONDO) [8] while standardizing phenotypic features using HPO terms. Consequently, it establishes a vital paradigm for rare disease phenomics and data sharing. Maintained jointly by the Global Alliance for Genomics and Health (GA4GH) core team and affiliated research groups, the Phenopacket Store advances the standardized storage, sharing, and analysis of clinical phenotypes.

In the present study, we aimed to investigate the feasibility of rare disease discrimination based on observed and excluded phenotypes. To this end, we first constructed a specialized dataset derived from the Phenopacket Store, and then applied nine machine learning methods to construct classification models for this task. Subsequently, feature selection was performed to assess the importance of the observed and excluded phenotypes using two machine learning methods. Based on further analysis, we obtained the conclusion. Our work is expected to establish a foundation for future development of highly accurate and practical diagnostic systems for rare diseases.

## Results

### Datasets, samples and features

In the early stages of diagnosis, rare diseases often present with only a limited set of clinical phenotypes, and whether these phenotypes are associated with a rare disease is unknown, rendering patients highly vulnerable to misdiagnosis. Consequently, to investigate whether machine learning can discriminate rare diseases from their controls based on these phenotypes became the primary motivation for our study. To achieve this, positive (rare disease) and negative (control) samples were required, respectively, to construct the training set. The positive samples were obtained from the Phenopacket Store (**Materials and Methods**). However, the Phenopacket Store lacks negative samples, which constituted the main bottleneck of the present study. To address this limitation, we generated a negative sample for each positive sample by using its excluded phenotypes as the observed phenotypes of the paired negative sample. Based on this strategy, we constructed a specialized dataset derived from the Phenopacket Store, designated as Rare1165. This dataset, encompassing 1,165 positive and 1,165 negative samples, was used for model training with cross-validation, whereas another dataset Rare1042, encompassing 1,042 additional positive samples, was constructed for independent testing. Although a total of 2,489 distinct phenotypes are represented across all samples (**Figure 1**), only 1,635 were selected as features to represent each sample, thereby reducing the risk of overfitting. Here, each sample corresponds to an individual clinical case, and each feature corresponds to a phenotype encoded as 0 or 1, where 1 indicates an observed state and 0 indicates an unobserved or unrecorded state.

**Figure 1.**
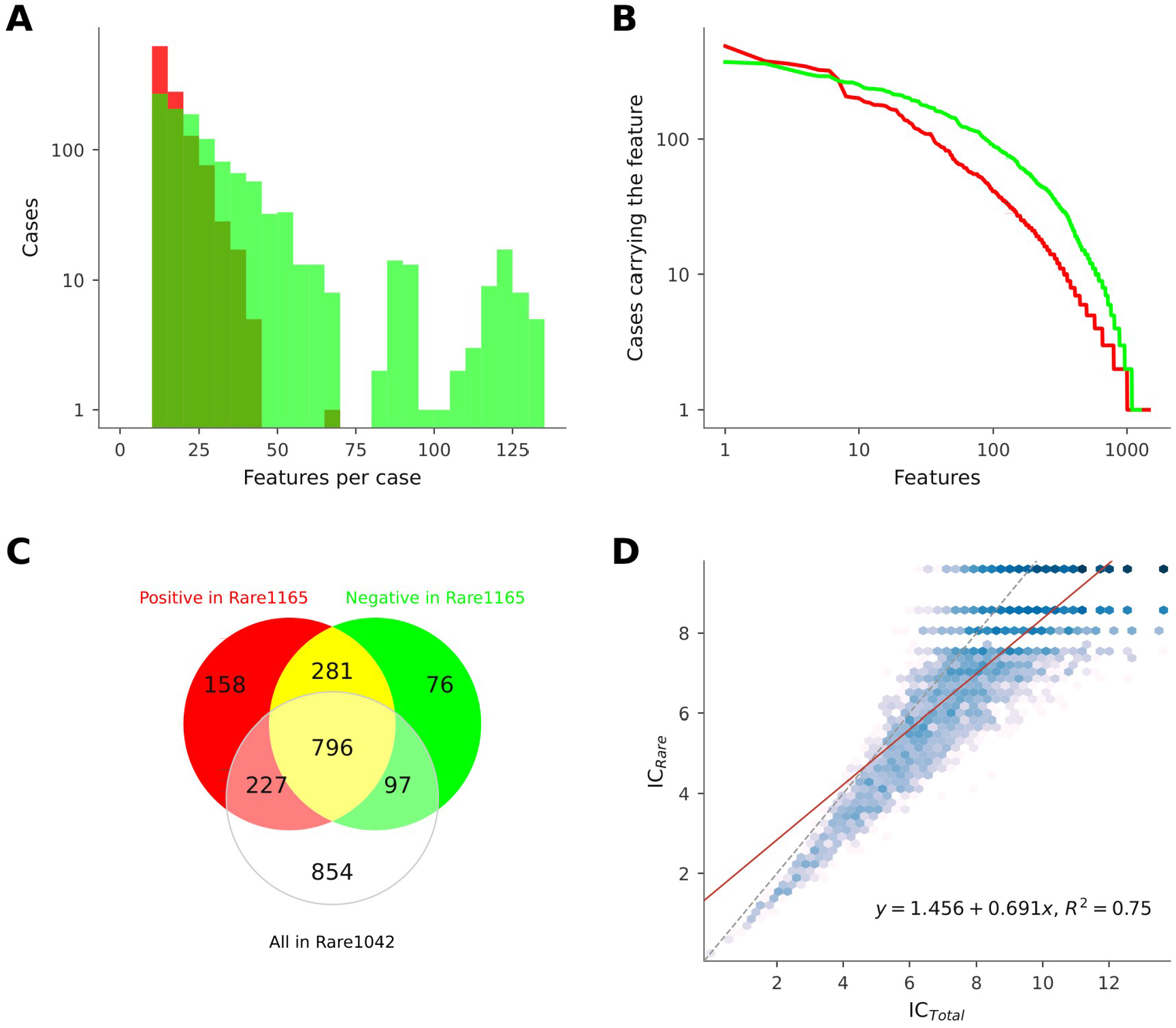
Overview of datasets, samples and features.

The training dataset Rare1165 consists of 1,165 positive (rare disease) samples and 1,165 negative samples, while the independent test dataset Rare1042 contains 1,042 additional positive samples (**Materials and Methods**). A total of 1,635 features were used to represent a sample. Here, each sample corresponds to an individual clinical case, and each feature corresponds to a phenotype with 0 and 1 encoding, where 1 indicates the observed state and 0 indicates the unobserved or unrecorded state. **Note** that the distribution of features per positive sample in Rare1042 (maximum: 46; minimum: 10; median: 13) is not shown in this figure. **A**. Distribution of features per positive sample in Rare1165 (maximum: 66; minimum: 10; median: 14) and per negative sample (maximum: 133; minimum: 10; median: 22); **B**. Distribution of positive sample coverage per feature in Rare1165 (maximum: 488; minimum: 0; median: 2) and negative sample coverage per feature (maximum: 372; minimum: 0; median: 4); **C**. Venn diagram illustrating the overlap of observed phenotypes across three groups: the 1,165 positive samples in Rare1165, the 1,165 negative samples in Rare1165, and the 1,042 positive samples in Rare1042. **D**. IC_Total_, was defined on the basis of its frequency across 12,823 total diseases in the HPO database v2025-11-24, whereas IC_Rare_, was defined across 780 rare diseases in Phenopacket Store v0.1.27. IC_Total_ and IC_Rare_ ranged from 1.64–13.65 (median 7.84) and 1.14–9.61 (median 6.61), respectively. The ratio of IC_Rare_ and IC_Total_ (RIC) ranged from 0.5183–1.4624 (median 0.8249).

For each HPO term, an important numerical metric—the information content (IC) [9]—can be used to quantify the specificity of its corresponding phenotype in clinical diagnosis and phenotype matching (**Materials and Methods**). In brief, the IC of an HPO term is positively correlated with the specificity of its corresponding phenotype: the higher the IC, the more specific the phenotype, and vice versa. In the present study, we defined two IC metrics derived from different data sources. The first, IC_Total_, was defined on the basis of its frequency across a total of 12,823 diseases in the HPO database v2025-11-24, whereas the second, IC_Rare_, was defined across 780 rare diseases in Phenopacket Store v0.1.27. Comparison of IC_Rare_ with IC_Total_ (**Figure 1D**) revealed a linear correlation between them (R^2^≈ 0.75), and the fitted slope (≈ 0.691) indicated that the values of IC_Rare_ were generally smaller than those of IC_Total_.

### Discriminating rare disease cases from their controls via machine learning

Using Rare1165 as the training dataset, we applied nine machine learning classification methods to discriminate rare disease cases from their controls using observed and excluded phenotypes as features. The nine methods to construct classification models included Support Vector Machine (SVM) [10], Random Forest (RF) [11], Generalized Boosted Model (GBM) [12], Recursive Partitioning and Regression Tree (RPART) [13], Artificial Neural Networks (ANN) [14], Partial Least Square (PLS) [15], K Nearest Neighbor (KNN) [16], Naive Bayes (NB) [17], and AdaBoost [18]. Receiver operating characteristic (ROC) curves of nine models were plotted (**Figure 2A**) and their performance (**Materials and Methods**) was evaluated under the standard 10-fold cross-validation framework using accuracy, Matthews Correlation Coefficient (MCC), and area under the ROC curve (AUC). The promising performance (**Table 1**) demonstrated the feasibility of classification on this dataset, indicating that machine learning can effectively discriminate rare disease cases from their controls using appropriately constructed features.

**Table 1.** Discriminating rare disease cases from their controls.

| Model | 10 cross-validation |  | Independent test |
| --- | --- | --- | --- |
|  | Accuracy | MCC | Accuracy |
| SVM-Linear | 0.9592 | 0.9185 | 0.7956 |
| RF | 0.9584 | 0.9174 | 0.9223 |
| GBM | 0.9455 | 0.8910 | 0.7898 |
| RPART | 0.8305 | 0.6626 | 0.5662 |
| ANN | 0.9562 | 0.9125 | 0.8455 |
| PLS | 0.9635 | 0.9271 | 0.8196 |
| KNN | 0.9562 | 0.9130 | 0.8935 |
| NB | 0.8614 | 0.7340 | 0.9779 |
| AdaBoost | 0.9528 | 0.9056 | 0.7466 |

Based on the Rare1165 dataset, the performance of nine classification models was evaluated using their optimal hyperparameters, which were determined through an exhaustive grid search with 10-fold cross-validation. Subsequently, nine models configured with these optimal settings were tested on an independent dataset Rare1042. The detailed search ranges and optimal configurations for all hyperparameters are provided in Supplementary File 1. Model performance was evaluated using accuracy and Matthews Correlation Coefficient (MCC). MCC could not be calculated for the independent test set, as it comprises exclusively positive samples. All classification models are trained and tested on the complete feature set, which includes a total of 1635 features.

**Figure 2.**
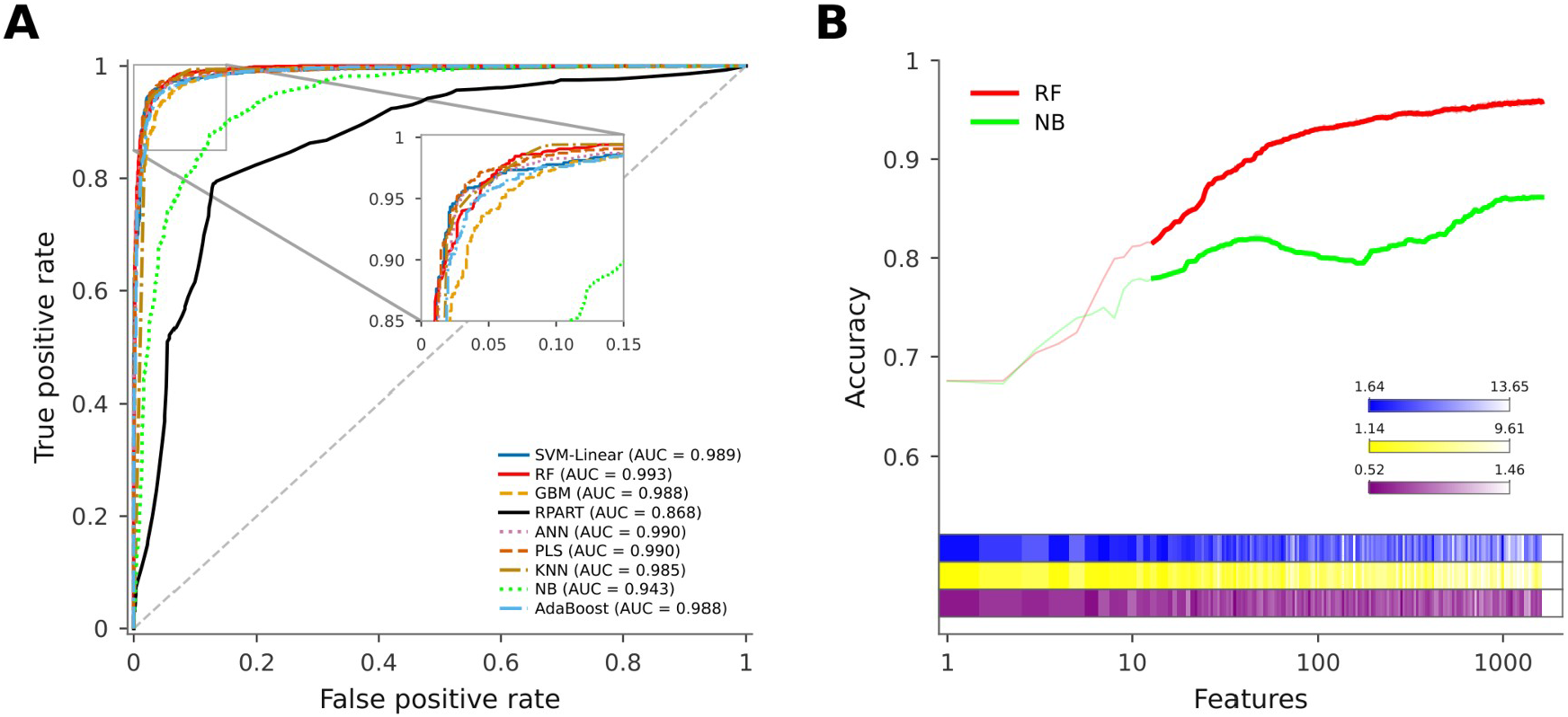
Model Performance on the Full Feature Set and Feature Subsets. **A.** Based on the Rare1165 dataset, the performance of nine models was evaluated using the optimal hyperparameters (**Supplementary File 2**) determined with the full feature set through an exhaustive grid search with 10-fold cross-validation; **B**. All feature-subset classification models were evaluated on the Rare1165 dataset with 10-fold cross-validation and hyperparameters that had been optimized on the full feature set under the same setting. The x-axis shows the top n features, where n ranges from 1 to 1,635. The heat maps (from top to bottom) in blue, yellow, purple colors represent the IC_Rare_, IC_Total_, and RIC values of features, respectively.

Independent testing on Rare1042 revealed that, among the nine evaluated models, NB and RF maintained their cross-validation performance, with NB improving slightly and RF declining only marginally (**Table 1**). These results suggest that RF and NB effectively captured latent relationships between features, rather than merely relied on sample-specific feature co-occurrences, thereby demonstrating a certain degree of generalizability. In contrast, the remaining models—including ANN, PLS, KNN, SVM-Linear, GBM, and AdaBoost—experienced varying degrees of performance decline, whereas RPART exhibited a substantial drop (**Table 1**). This performance decline was most likely due to the low feature overlap between Rare1042 and Rare1165. Although 1,120 phenotype terms were shared across the two datasets, the 854 phenotype terms unique to Rare1042 were excluded from the feature set used for model training (**Figure 1C**).

### Feature selection with further analysis

The performance decline observed in the independent test underscored the critical influence of feature representation. We therefore performed feature selection to assess feature importance, *i*.*e*., the contribution of each feature to the model’s classification performance (**Materials and Methods**). Only the RF and NB classification models were employed for feature selection (**Figure 2B**), as both maintained cross-validation performance in the independent test. The random forest importance spectrum (RF-IS) method [19] was used to generate the feature importance ranking, which was validated by cumulative feature performance curves constructed with RF models, while the univariate filtering feature selection (UFFS) method [20] based on the chi-square test was also used and its ranking results were independently validated by cumulative feature performance curves constructed with NB models. As shown in Figure 2B, the RF model with the top 43 features achieved an accuracy of 0.9073, whereas its highest accuracy of 0.9622 was obtained using 1,398 features. Similarly, the NB model with the top 43 features achieved 0.8206, while its highest accuracy of 0.8730 was obtained using 933 features.

As illustrated in **Figure 2B**, classification performance exhibited a clear monotonic dependence on the number of included features. The performance plateau after a certain growth stage indicated that a large fraction of the initially used features are redundant, or have only weak or no associations with rare diseases. The redundancy originates from the inherent structure of HPO: as a directed acyclic graph (DAG), HPO organizes its terms in a hierarchical manner with substantial semantic redundancy, which is particularly reflected in the cumulative feature performance curve of NB models (**Figure 2B**). The structure of HPO enables full semantic coverage of heterogeneous phenotype documentation, which addresses the common scenario where different clinicians describe the same underlying clinical phenotype using different HPO terms. Unexpectedly, we observed that features with higher importance rankings tend to exhibit lower IC_Rare_ and IC_Total_ values. Notably, this trend was even more pronounced for the ratio between a feature’s IC_Rare_ and IC_Total_, termed relative information content (RIC): RIC decreased more steeply with increasing importance ranking than either IC_Rare_ or IC_Total_ alone (**Figure 2B**).

Base on the above observation, we further empirically demonstrated that features with lower RIC values yield better classification performance across feature-subsets of varying sizes. For example, using only a single phenotype, HP:0001263 (**Table 2**), the RF and NB models achieved the best classification accuracy of 0.6755 on the Rare1165 dataset. The RIC of HP:0001263 is only 0.6098. As another example, among all 19,600 three-feature subsets selected from top 50 features, the subset consisting of HP:0001263, HP:0031936, and HP:0001270 yielded the best classification accuracy of 0.7309 with the RF model. Notably, all three features had low RIC values (0.6098, 0.5654, and 0.5404, respectively), further supporting our finding (**Table 3**). This systematic trend can be explained by the formulation of IC, which is defined under the well-established true-path rule (**Materials and Methods**). Under this rule, the IC values of parent nodes in the HPO DAG are systematically lower than those of their child nodes. However, a certain proportion of subsets with low RIC values still achieved poor classification performance, caused by the insufficient occurrence counts in the training dataset (**Table 3**). Consequently, the IC metric alone is not suitable for feature construction and selection in rare disease discrimination.

**Table 2.** Top 10 selected features with information content.

| Feature* | Term | IC<sub>Total</sub> | IC<sub>Rare</sub> | RIC | Positive | Negative |
| --- | --- | --- | --- | --- | --- | --- |
| HP:0001263 | Global developmental delay | 2.2061 | 1.3452 | 0.6098 | 488 | 79 |
| HP:0001629 | Ventricular septal defect | 4.2977 | 3.0682 | 0.7139 | 57 | 362 |
| HP:0000341 | Narrow forehead | 5.8072 | 3.8260 | 0.6588 | 41 | 226 |
| HP:0001252 | Hypotonia | 2.2552 | 1.6243 | 0.7202 | 377 | 155 |
| HP:0001647 | Bicuspid aortic valve | 6.5275 | 4.7004 | 0.7201 | 16 | 202 |
| HP:0000750 | Delayed speech and language development | 3.1616 | 1.8129 | 0.5734 | 321 | 55 |
| HP:0001249 | Intellectual disability | 1.9521 | 1.6130 | 0.8263 | 361 | 60 |
| HP:0000316 | Hypertelorism | 3.3645 | 2.3035 | 0.6846 | 325 | 262 |
| HP:0000486 | Strabismus | 3.2014 | 2.2236 | 0.6946 | 178 | 327 |
| HP:0000365 | Hearing impairment | 2.6450 | 2.3594 | 0.8920 | 109 | 270 |

\*Features were ranked by the Random Forest-Importance Score (RF-IS) method. IC<sub>Total</sub> was defined on the basis of its frequency across a total of 12,823 diseases in the HPO database v2025-11-24, whereas IC<sub>Rare</sub> was defined across 780 rare diseases in Phenopacket Store v0.1.27. Relative information content (RIC) is formally defined as the ratio between IC<sub>Rare</sub> and IC<sub>Total</sub>. Column 7 shows the occurrence of this term in positive samples from the Rare1165 dataset, and Column 8 shows that in negative samples.

**Table 3.** Performance of classification models using feature subsets.

| Subset | Features used in RF models |  |  | Accuracy | MCC |
| --- | --- | --- | --- | --- | --- |
| Top 10 | The top 10 RF-IS–selected features (Table 2) |  |  | 0.8206 | 0.6449 |
| Top 6/10 | The 1st, 2nd, 3rd, 5th, 6th and 9th of Top 10 |  |  | 0.7579 | 0.5421 |
| Top 4/10 | The 4th, 7th, 8th and 10th of Top 10 |  |  | 0.7060 | 0.4129 |
| Group1 | HP:0001263 | HP:0000750 | HP:0001249 | 0.7369 | 0.4922 |
| Group2 | HP:0001263 | HP:0031936 | HP:0001270 | 0.7309 | 0.4809 |
| Group3 | HP:0001629 | HP:0001762 | HP:0000582 | 0.7300 | 0.4854 |
| Group4 | HP:0001762 | HP:0001631 | HP:0000490 | 0.7292 | 0.4771 |
| Group5 | HP:0001629 | HP:0100021 | HP:0001762 | 0.7288 | 0.4937 |
| Group6 | HP:0001263 | HP:0000750 | HP:0011344 | 0.7279 | 0.4804 |
| Group7 | HP:0001263 | HP:0000750 | HP:0034391 | 0.7279 | 0.4822 |
| Group8 | HP:0001263 | HP:0000750 | HP:0031936 | 0.7258 | 0.4712 |
| Features used in NB models |  |  |  |  |  |
| Top 10 | The top 10 RF-IS–selected features (Table 2) |  |  | 0.7687 | 0.5565 |
| Top 6/10 | The 1st, 2nd, 3rd, 5th, 6th and 9th of Top 10 |  |  | 0.7489 | 0.5201 |
| Top 4/10 | The 4th, 7th, 8th and 10th of Top 10 |  |  | 0.6687 | 0.3630 |

The second column lists all features in each subset, which are presented either as Human Phenotype Ontology (HPO) IDs or as their corresponding rankings derived from the Random Forest-Importance Score (RF-IS) procedure on the full feature set. All classification models constructed on feature-subsets using the Random Forest (RF) or Naive Bayes (NB) methods were evaluated on the Rare1165 dataset with 10-fold cross-validation and hyperparameters that had been optimized on the full feature set under the same setting. Model performance was evaluated using accuracy and Matthews Correlation Coefficient (MCC). Among all 19,600 three-feature subsets selected from top 50 features, Groups 1 to 8 yielded the top 8 classification accuracies with the RF model.

Among the top 10 RF-IS–selected features (**Table 2**), four features ranked 4th, 7th, 8th, and 10th, were mapped to high-level terms in the HPO DAG (**Figure 3A**), while the remaining six features ranked 1st, 2nd, 3rd, 5th, 6th, and 9th, were mapped to relatively low-level terms. The four features yielded poorer classification performance than the remaining six (**Table 3**). Then, we ruled out the possibility that this systematic trend originated from the Phenopacket Store’s phenotype annotation rule, which differs substantially from the true-path rule in IC formulation. Specifically, when the Phenopacket Store annotates each rare disease case with a designated HPO term, none of its ancestor or descendant terms in the HPO DAG will be tagged simultaneously. Accordingly, the occurrence counting of any HPO term across rare disease cases is independent of that of its ancestor or descendant terms. Based on these results, we concluded that phenotypes with high specificities contribute more to rare disease discrimination using machine learning methods. Further quantitative analysis revealed that the terms corresponding to the top 10 features have significantly higher occurrence counts than their respective ancestor or descendant terms (**Figure 3BC**). Additionally, we found that among the top 10 features, five were initially selected from observed phenotypes and the other five from excluded phenotypes, suggesting that observed and excluded phenotypes make comparable contributions to rare disease discrimination. In summary, we conclude that the two factors, specificity and occurrence count, are critically important for phenotype selection in rare disease discrimination, and comparable importance should be attached to both observed and excluded phenotypes.

**Figure 3.**
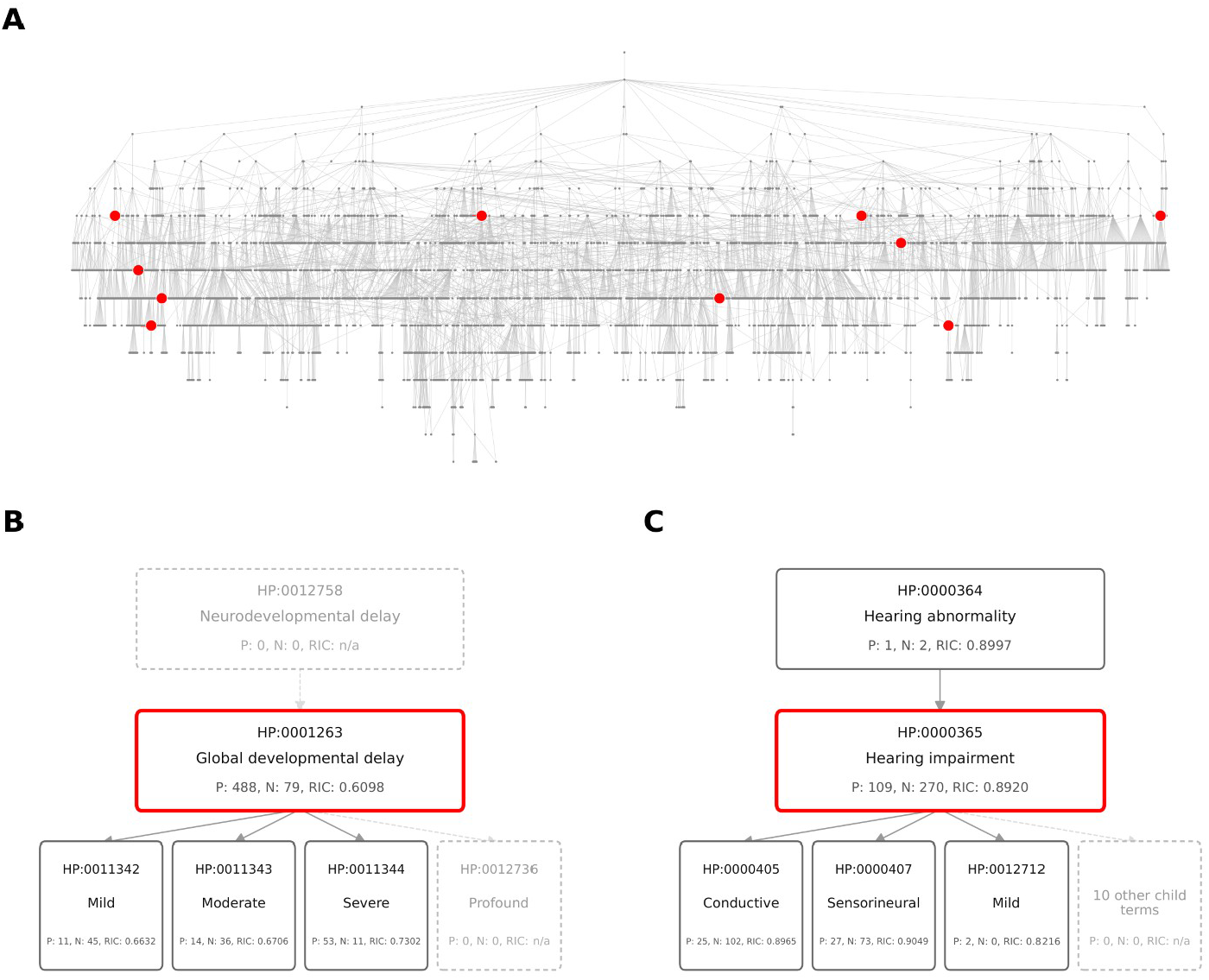
Occurrence count, importance and specificity of HPO terms. **A.** The DAG displays the full feature set corresponding to 1,635 HPO terms and their inherent ancestor–descendant hierarchical associations, with all nodes rendered in grey. The top 10 features selected by RF-IS (**Table 2**) are highlighted in red, arranged sequentially from left to right and top to bottom as HP:0000316, HP:0001252, HP:0001249, HP:0000365, HP:0001263, HP:0000486, HP:0001629, HP:0000341, HP:0001647, and HP:0000750. **B**. P and N represent the occurrence counts of this term in positive samples and negative samples in the Rare1165 dataset, respectively and RIC represents relative information content. **C**. HP:0001263 and HP:0000365 were ranked as top 1 and 10 (Table 2).

## Materials and methods

### Data acquisition, dataset construction and sample encoding

The initial dataset was retrieved from Phenopacket Store v0.1.27 (https://github.com/monarch-initiative/phenopacketstore/releases/download/0.1.27/all_phenopackets.zip), comprising a total of 10,377 samples, each corresponding to an individual clinical case. The data filtering pipeline was executed as follows: first, cases with fewer than 10 observed phenotypes were excluded, leaving 3,760; second, only cases assigned an ORPHA code were retained, resulting in 2,666 cases. The 2,666 cases were subsequently grouped by their ORPHA codes into 278 disease categories, from which categories with fewer than 5 samples were removed, reducing the count to 119 categories. Finally, two non-specific categories—ORPHA:528084 and ORPHA:442835—were excluded. Ultimately, a total of 2,207 cases spanning 117 categories were retained.

Of the 2,207 cases, 1,165 possessing more than 10 excluded phenotypes were selected as positive samples. For each positive sample, a negative sample was generated by using its excluded phenotypes as the observed phenotypes of the paired negative sample. This paired design yielded 1,165 positive-negative sample pairs, which were subsequently used to construct the training dataset, designated as Rare1165. The remaining 1,042 unpaired positive samples were used to construct the independent test dataset, designated as Rare1042. The Rare1165 and Rare1042 datasets are available in **Supplementary File 2**.

In Rare1165, the positive samples encompassed 1,462 phenotypes, whereas the negative samples encompassed 1,250 phenotypes (**Figure 1**). Specifically, 1,077 phenotypes were shared between the two groups, while 385 and 173 phenotypes were unique to the positive and negative groups, respectively. The union of these phenotypes—1,077 + 385 + 173 = 1,635 in total—was adopted as the feature set to represent each individual sample. Between Rare1042 and Rare1165, 1,120 phenotypes were shared, while 854 and 515 were unique to Rare1042 and Rare1165, respectively; notably, the 854 phenotypes unique to Rare1042 were excluded from the feature set. Each sample was then represented by binary encoding (1/0), where 1 indicates the observed state and 0 indicates the unobserved or unrecorded state.

### Model training, tuning and feature selection using machine learning

SVM, RF, GBM, RPART, ANN, PLS, KNN, NB, and AdaBoost were briefly described in our previous study [21]. In the present study, these methods were used to construct classification models on the Rare1165 dataset with their optimal hyperparameters, which were determined through exhaustive grid search with 10-fold cross-validation. All model training and tuning procedures were implemented using the caret package [22] v6.0-94 on R 4.3.3. After evaluating SVM methods with two distinct kernel functions, SVM with a linear kernel (SVM-Linear) was selected for further analysis because it yielded performance comparable to SVM with a radial kernel (SVM-Radial), while being substantially more computationally efficient.

RF-IS, a RF-based method for feature selection, was implemented using the scikit-learn package [23] v1.8.0 on Python 3.12.3. RF-IS incorporates two evaluation metrics: Mean Decrease Accuracy (MDA) and Mean Decrease Impurity (MDI), the latter of which is calculated based on the Gini index. In the present study, the MDI metric was selected to score feature importance and all features were subsequently ranked in descending order based on their computed MDI scores. Based on the class RandomForestClassifier, an in-house Python script was developed to perform Recursive Feature Elimination (RFE): in each iteration, the bottom 20% of features were discarded and their ranks recorded, while the remaining features were re-ranked according to their MDI scores. This procedure was repeated until termination, yielding a complete descending ranking of feature importance.

In the present study, the UFFS method is based on the chi-square test to score feature importance, implemented using the scikit-learn package v1.8.0 on Python 3.12.3. Based on the class SelectKBest, an in-house Python script was developed to extract a complete descending importance ranking for all features. To validate the feature importance ranking by RF-IS and UFFS, we constructed cumulative feature performance curves using the RF and NB classification models, respectively. These curves depict classification accuracy (y-axis) as a function of the number of top features included in the model (x-axis).

### Performance evaluation, cross validation and information content

Model performance was evaluated using accuracy, MCC, and AUC. These measures were described in detail in our previous study [24] and calculated using in-house R scripts. IC is used to reflect specificity of a HPO term. In the formula IC(*t*)=−log_2_*f*(*t*), t denotes an HPO term, and f(t) refers to its frequency, defined as the proportion of diseases annotated with this term out of the total number of diseases. The formulation of IC is defined under the well-established true-path rule. Under this rule, if a term is annotated to a clinical case, all its ancestor terms along the directed path to the HPO root node are implicitly annotated as well. Accordingly, every raw phenotype annotation is first propagated upwards across the entire DAG hierarchy before the term frequency counting step for IC calculation. This inherent propagation mechanism ensures that the IC of terms increases progressively as one traverses from the ontology root node toward its more specific descendant terms.

## Conclusion

The present study revealed that the two factors, specificity and occurrence count, are important for phenotype selection in rare disease discrimination, and comparable importance should be attached to both observed and excluded phenotypes, during feature construction. However, these two factors are still, to a certain extent, subject to the phenotype documentation practices of clinicians for the target rare diseases. Therefore, the design of a diagnostic system based on HPO terminology with built-in recommendation functionality would facilitate timely diagnosis of rare diseases.

All results and conclusions of the present study were derived from the currently available datasets, so the inherent limitations introduced by the datasets themselves cannot be completely avoided. Potential issues such as incomplete coverage of rare disease types or biased documentation of phenotypes may lead to deviation or even errors in the analytical results and conclusions. For instance, the absence of certain phenotypes in rare disease records may not stem from the fact that these features were not actually exhibited in patients, but from the selective recording bias, where clinicians did not document these features as they consider them to have low clinical relevance to the target rare disease. Despite the above limitations, the preliminary exploration in the present study still points to clear directions for subsequent research, and provides new insights to advance the early diagnosis of rare diseases.

## Authors’ contributions

Shan Gao conceived the project. Shan Gao and Guangyou Duan supervised this study. Yanting Guo and Cihan Ruan performed programming. Yang Yao and Yiyao Zhang downloaded, managed and processed the data. Shan Gao drafted the main manuscript text. Shan Gao and Guangyou Duan revised the manuscript.

### Acknowledgments

We sincerely thank Xin Li from College of Life Sciences at Nankai University. This manuscript was submitted to xx as a preprint on Sep. 15th, 2026.

